# From pathogen to commensal to probiotic: modification of *Microbacterium nematophilum-C. elegans* interaction during chronic infection by the absence of host insulin signalling

**DOI:** 10.1101/2020.04.01.020933

**Authors:** Maria Gravato-Nobre, Jonathan Hodgkin, Petros Ligoxygakis

## Abstract

The nematode worm *Caenorhabditis elegans* depends on microbes in decaying vegetation as its food source. To survive in an environment rich in opportunistic pathogens, *C. elegans* has evolved an epithelial defence system where surface-exposed tissues such as epidermis, pharynx, intestine, vulva and hindgut have the capacity of eliciting appropriate immune defences to acute gut infection. However, it is unclear how the worm responds to chronic intestinal infections. To this end, we have surveyed *C. elegans* mutants that are involved in inflammation, immunity and longevity to find their phenotypes during chronic infection. Worms that grew in a monoculture of the natural pathogen *Microbacterium nematophilum* (CBX102 strain) had a reduced lifespan and health span. This was independent of intestinal colonisation as both CBX102 and the derived avirulent strain UV336 were early persistent colonisers. In contrast, long-lived *daf-2* mutants were resistant to chronic infection, showing reduced colonisation and a higher age-dependent vigour. In fact, UV336 acted as a probiotic in *daf-2*, showing a lifespan extension beyond OP50, the *E. coli* strain used for laboratory *C. elegans* culture. Longevity and vigour of *daf-2* mutants growing on CBX102 was dependent on the FOXO orthologue DAF-16. Since the DAF-2/DAF-16 axis is present in most metazoans this suggests an evolutionary conserved host mechanism to modify a pathogen to a commensal.

## INTRODUCTION

Animal epithelia from hydra to humans possess innate mechanisms that sense pathogenic and toxic insults and transmit non-self/danger recognition signals to activate appropriate defences (Zasloff 2002; Bartlett 2008; Augustin *et al*, 2012). The efficacy of these systems determines whether microbial populations can be controlled, and thus organismal homeostasis maintained. *C. elegans* is a bacterial feeder that spends much of its life in decomposing vegetable matter and depends on microbes as its food source (Frezal and Felix 2015). These microbes are ground by the pharynx before they subsequently enter the gut. To survive in an environment rich in potentially damaging microorganisms, *C. elegans* has evolved an epithelial defence system coupled with the ability to discriminate between pathogenic vs. edible bacteria (reviewed in Kim and Ewbank, 2018).

Important antimicrobial molecules participating in these defences include a group of proteins called invertebrate lysozymes (ILYS) and in particular ILYS-3, which is expressed in both the pharynx and the intestine (O’Rourke *et al*, 2006). ILYS-3 (invertebrate-specific but related to human epithelial antimicrobial peptides) contributes to the digestion of the large amount of peptidoglycan fragments generated by the worm’s bacterial diet (either pathogenic or non-pathogenic) (Gravato-Nobre *et al*, 2016). Loss of *ilys-3* results in colonization of undigested bacteria from day 1 of adulthood in contrast to wild type worms (Gravato-Nobre *et al*, 2016). The latter only display colonization at very late stages of their life (Gravato-Nobre *et al*, 2016). Increased bacterial colonization in *ilys-3* mutants leads to a significant lifespan reduction (Gravato-Nobre *et al*, 2016).

The isolation of natural bacterial pathogens of *C. elegans* has permitted a glimpse of the defence mechanisms employed by the worm as well as the host-pathogen interactions triggering such mechanisms (see Hodgkin *et al*, 2000; Nicholas and Hodgkin 2004; Hodgkin *et al* 2013). One such pathogen is *Microbacterium nematophilum* (Hodgkin *et al*, 2000). This Gram-positive bacterium adheres to the rectal and anal cuticle, *Microbacterium nematophilum* (Hodgkin *et al*, 2000) and induces inflammation, anal-region infection and tail swelling (Parsons and Cipollo, 2014). Despite the fact that the most obvious response to infection is rectal colonization and the induction of inflammation in the rectal tissues, this bacterium also establishes itself in the gut of the worm. This makes it a good system to investigate effects that occur in the digestive tract associated with long-term gut colonization. In particular, to identify how longevity and health of the organism can be achieved in the face of chronic intestinal infection.

To explore this question, we tested *C. elegans* mutants induced by chemical mutagenesis or targeted deletion in signalling pathways known to be involved in immunity to *M. nematophilum* infection and/or *C. elegans* longevity. Culturing worms on the pathogenic *M. nematophilum* strain CBX102 was able to separate estimated survival probabilities into four categories in relation to *ilys-3* and wild type worms and identified *daf-2* as long-lived in conditions of chronic infection. Bacterial colonisation of CBX102 in wild type (N2) worms was increased compared to the laboratory *E. coli* strain OP50. However, colonisation in N2 *per se* was not the reason for pathogenesis as the non-virulent *M. nematophilum* strain UV336 did not curtail lifespan despite being able to colonise at the same levels as CBX102. Nevertheless, *daf-2* worms were healthier and had reduced colonisation compared to normal worms. *daf-2* health and longevity on CBX102 involved the canonical insulin signalling pathway and were thus dependent on the FOXO orthologue *daf-16*, like many other *daf-2-*mediated effects. Finally, the non-pathogenic UV336 was able to support an extended lifespan for *daf-2* even compared to OP50. These results indicate the complex and strain-specific interactions between intestinal bacteria and their host.

## RESULTS

### Chronic Gastrointestinal Infection (CGI) curtails lifespan, reduces health and accelerates ageing in N2 worms

In our experimental set-up, *C. elegans* develops, feeds and ages in a monoculture of *M. nematophilum*, having the same immune pressure from birth. Compared to standard laboratory food (*E. coli* strain OP50), the pathogenic *M. nematophilum* strain CBX102 accelerated age-dependent bacterial colonization (see below). CGI reduced host lifespan (Fig. 1A) and health measured by vigour of movement in liquid assays (Fig. 1B). The avirulent *M. nematophilum* UV336 strain (derived from CBX102 by UV mutagenesis, Akimkina *et al*, 2006), had the same level of age-dependent bacterial colonisation as CBX102 (Fig. 1C) but in contrast to the latter, presented no negative impact on median lifespan (Fig. 1A) or health span (Fig. 1B) both of which were largely comparable to OP50. In this context, two strains of the same species behaved one as a pathogen (CBX102) and one as a commensal (UV336). Moreover, CBX102 accelerated mitochondrial fragmentation (Fig S1), a sign of age-dependent stress in worms (Han *et al*, 2017).

**FIGURE 1.**
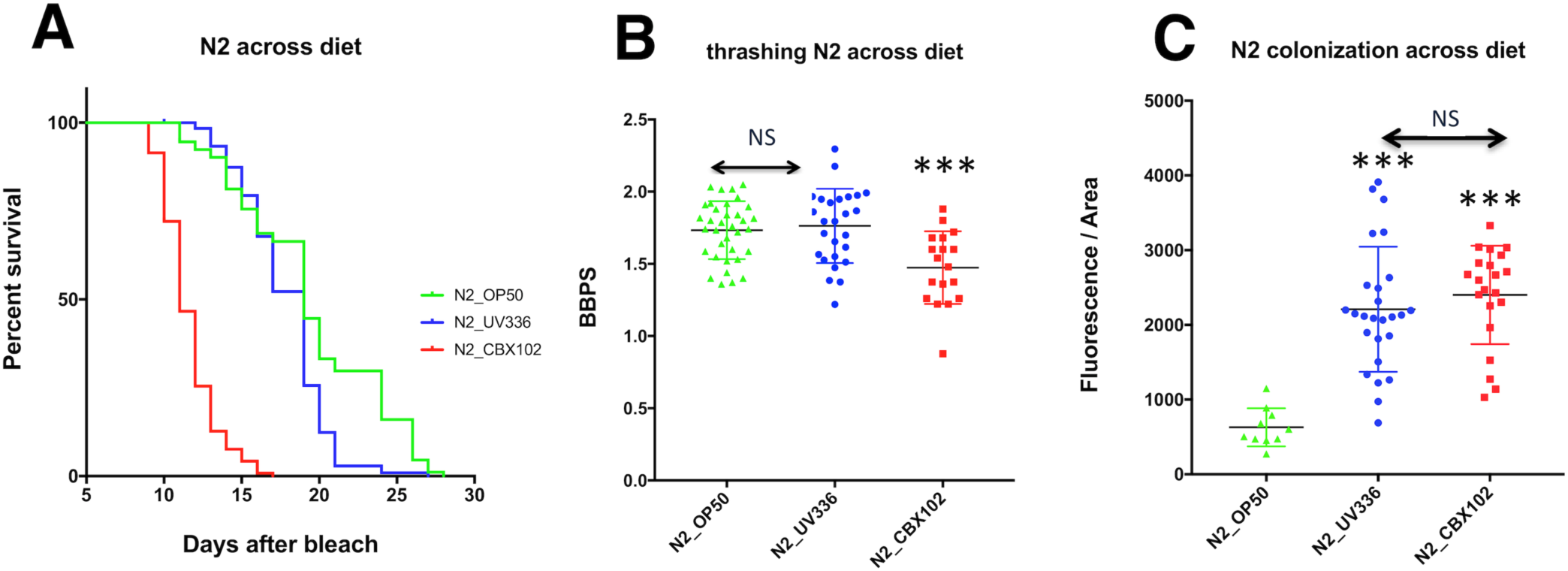
Lifespan, health and bacterial colonisation of the reference strain N2 in UV336 vs. CBX102. **(A)** Lifespan analysis at 25°C showing that CBX102 (red) significantly reduced average survival calculated using the Mantel-Cox log-rank test, 95% Confidence Interval (CI) compared to UV336 and OP50. The latter strains were statistically indistinguishable (NS). **(B)** Rigorous movement (thrashing) of animals grown on OP50, UV336 or CBX102 as a proxy for health was calculated as the number of body bends per second (BBPS). Tukey’s multiple comparisons with one-way ANOVA test was performed. Worms on CBX102 were significantly less mobile than on OP50 or UV336. These were again, statistically indistinguishable (NS). **(C)** Shown are distributions for the fluorescence intensity of SYTO13 in the intestine of animals on OP50 (*E. coli*), UV336 (*M. nematophilum*, non-inflammatory strain) and CBX102 (*M. nematophilum* pathogenic strain) at 25°C. Asterisks indicate the results of Two-Tukey’s multiple comparisons one-way ANOVA tests, 99% CI. All panels: ***p<0.0001, NS=non-significant and n=25 animals/treatments/group. Results are from 3 independent experiments.

### CGI defines four lifespan groups of *C. elegans* mutants

it is generally accepted that CGI has four components (see Stecher *et al*, 2015): These are 1) the infectious agent inducing the disease, 2) host genetics that will influence mucosal barrier function and the level of pro or anti-inflammatory responses, 3) the intestinal commensal microbiota that can enhance the disease when its composition change and 4) diet, which interacts with all other components as well as host metabolism. Negative interaction of these factors can abolish normal intestinal barrier function leading to constant mucosal inflammation and reduced health span and life expectancy (Finch 2010). In contrast, non-inflammatory microbiota can lead to extension of lifespan and health span (Hooper and Gordon 2001). It is evident that interactions of the above 4 components generate a complex set of conditions, which makes it hard to untangle the layers of chronic disease and arrive at causality. However, causality will ultimately define future therapeutic targets for health span extension.

In our simplified system, the nematode worm develops, feeds and ages in a bacterial monoculture. This means that food=microbiota=pathogen (or commensal) depending on the choice of bacterium. This condition ensures the ability to modify host genetics *in vivo* by keeping all other parameters important for CGIs in tight control. Although when the pathogen changes so will the function of diet and microbiota, the system enables in principle to find each time, the host genes that interact with a specific bacterium. With this in mind, we have tested whether mutants induced by chemical mutagenesis in signalling pathways known to be involved in immunity to *M. nematophilum* infection and/or *C. elegans* longevity, modulated intestinal colonization, lifespan and health span across the life course. All strains were cultured from eggs in pure CBX102 and tested for bacterial colonization. *The purpose was to find mutants that could outlive N2 under CGI while retaining their health*.

The mutants tested were of genes involved in evolutionary conserved innate immune response pathways against *M. nematophilum* and/or bacterial infection (e.g. the p38 MAPK pathway components *sek-1, nsy-1, pmk-1, kgb-1*; TGF-β with *dbl-1;* ERK with *sur-2*), cuticle properties (*sqt*-*3*), bacterial killing (the lysozyme-encoding *lys-3* and *lys-7*), pharyngeal-defective with enhanced bacterial colonisation of the intestine (*phm-2*), stress-specific regulators (*hsf-1*), apoptosis (the *p53* homologue *cep-1* and *ced-1*) and lifespan determinants (*hif-1, vhl-1, age-1, eat-2, cik-1, daf-2*). CGI separated the mutants tested into four categories: A) Those whose lifespan was shorter than *ilys-3* mutants (Fig 2A); B) those that had lifespan comparable to *ilys-3* (Fig. 2B); C) those with life expectancy comparable to N2 (Fig 2C); and D) those that had an increased lifespan compared to N2 (Fig 2D). Most of the time (but not always) bacterial colonisation negatively correlated with lifespan (Fig S2). Table S1 has a summary of alleles used categorised in the four groups as above (A-D) and includes lifespan, health (vigorous movement) and bacterial colonisation results along with extracted p-values for statistical significance. An exception was the *clk-1* mutant, which showed enhanced bacterial colonisation and yet lived longer but without showing any movement (data not shown).

**FIGURE 2.**
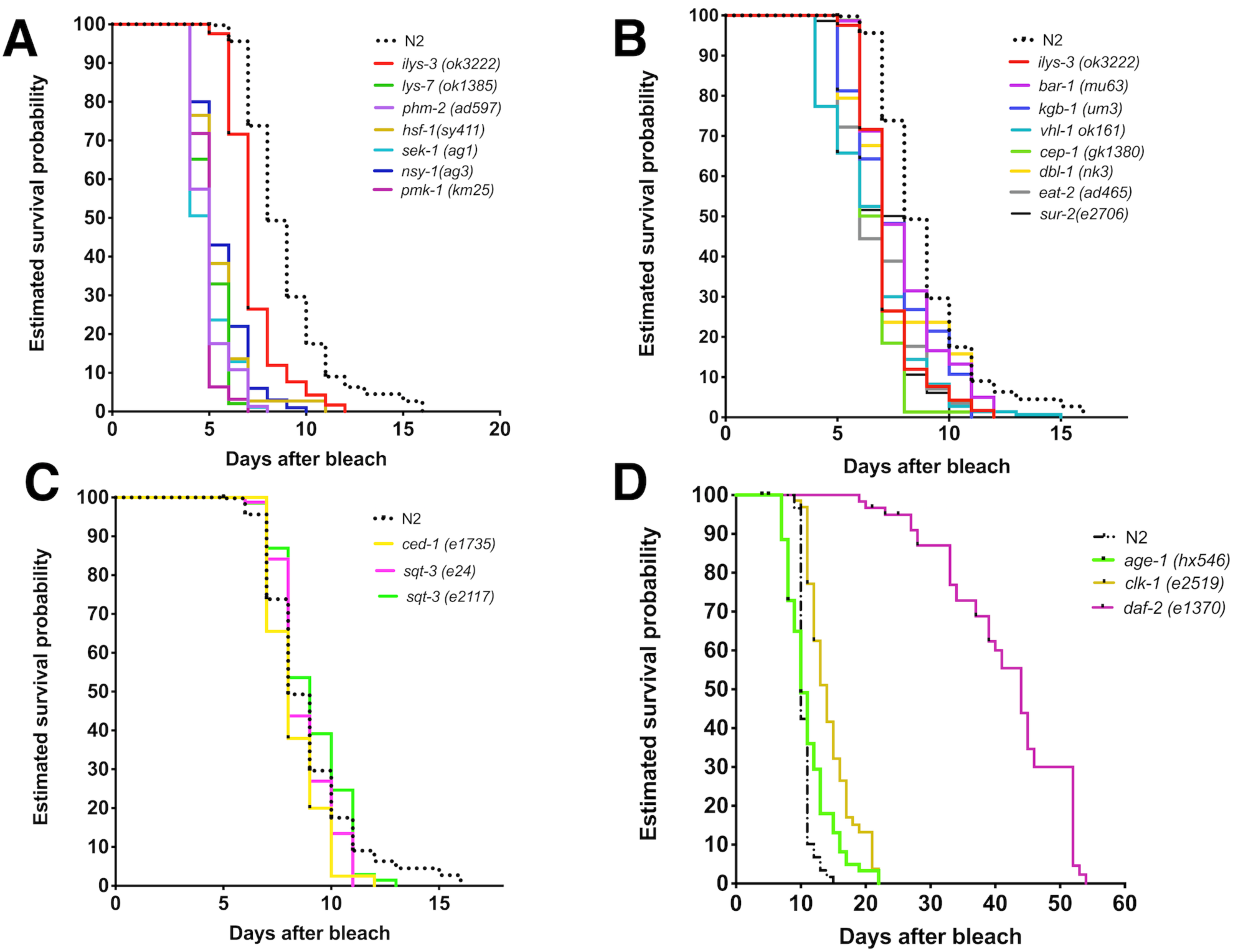
Lifespan of *C. elegans* mutants define 4 groups on the pathogenic *M. nematophilum* strain CBX102. **(A)** Mutations that significantly shorten the lifespan compared to *ilys-3*. TD50 =5 days. **(B)** Mutations that shorten the lifespan to the same degree as *ilys-3*. **(C)** Mutations with the same TD50 as N2 (8-9 days). **(D)** Mutations that extended the average survival compared to N2 (e.g. *daf-2*=44 days). N=100 animals/curve.

### *daf-2* mutant is long-lived and healthier than N2 under CGI

From the mutants tested, only one mutant in the insulin receptor *daf-2* was found to be living longer under CGI (Fig. 2D). This confirmed and extended observations for *daf-2* longevity in OP50 (Kenyon *et al*, 1993) as well as acute infections by *S. aureus, P. aeruginosa* or *E. faecalis* (Garsin *et al*, 2003) and *Salmonella typhimurium* (Portal-Celhay *et al*, 2012). Bacterial colonisation of *daf-2* was reduced compared to N2 (Fig. S2). It was also reduced compared to other normally long-lived mutants such as *age-1* (Fig S2). The latter is long-lived on OP50 (Friedman and Johnson, 1988) but had lifespan indistinguishable to N2 on CBX102. (Fig 2D).

Despite the adverse effects of CBX102 on N2 lifespan (when compared to OP50), N2 median lifespan on UV336 vs. OP50 was statistically indistinguishable (Fig. 3A). The survival pattern of *daf-2* mutants on CBX102 was statistically comparable to that of *daf-2* on *E. coli* OP50 (Fig. 3B). Compared to N2 on CBX102, *daf-2* worms were still longer-lived (compare Fig. 3A and 3B). Notably, *daf-2* lifespan was extended on UV336 compared to *daf-2* on CBX102 even beyond the TD_50_ and maximum lifespan limits defined by OP50 (Fig 3B). This boosting effect on lifespan by UV336 over and above OP50 was not observed in N2 (Fig. 3A). This result made the effects of *M. nematophilum* host genotype-specific and identified *daf-2* as a host genotype where UV336 acted as a probiotic. Moreover, this showed that the genotype of the host can modify the effect of a bacterial strain and this interaction determines lifespan (see discussion below).

**FIGURE 3.**
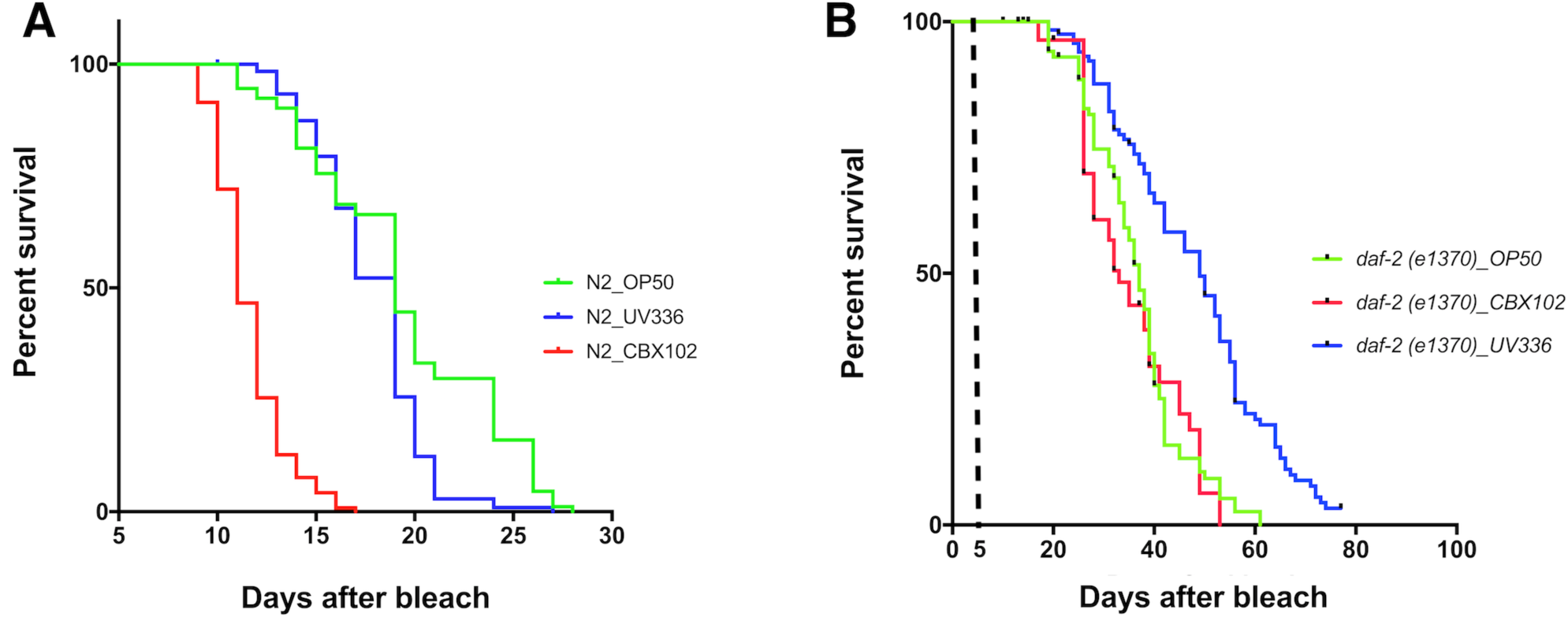
The *daf-2* mutant modifies the effects on lifespan of *M. nematophilum* strains. **(A)** Lifespan of N2 on *M. nematophilum* CBX102 under CGI was significantly reduced (TD_50_=11 days) when compared to both the derived *M. nematophilum* UV336 strain as well as *E. coli* OP50 that produced identical TD_50_ (19 days). **(B)** Lifespan of *daf-2* on *M. nematophilum* CBX102 under CGI (TD_50_=33) was statistically indistinguishable (p=0.4531) to OP50 (TD_50_=36). In contrast, lifespan on UV336 was significantly (p<0.0001) increased (TD_50_=49). For experiments involving the temperature sensitive *daf-2*, lifespan assays started at day 0 when animals were age-synchronized by bleach. Embryos were then left at 15^0^C on the appropriate bacterial diet till day 5. Day 5 marks the L4 to adult transition and time when plates were transferred to 25^0^C.

**FIGURE 4.**
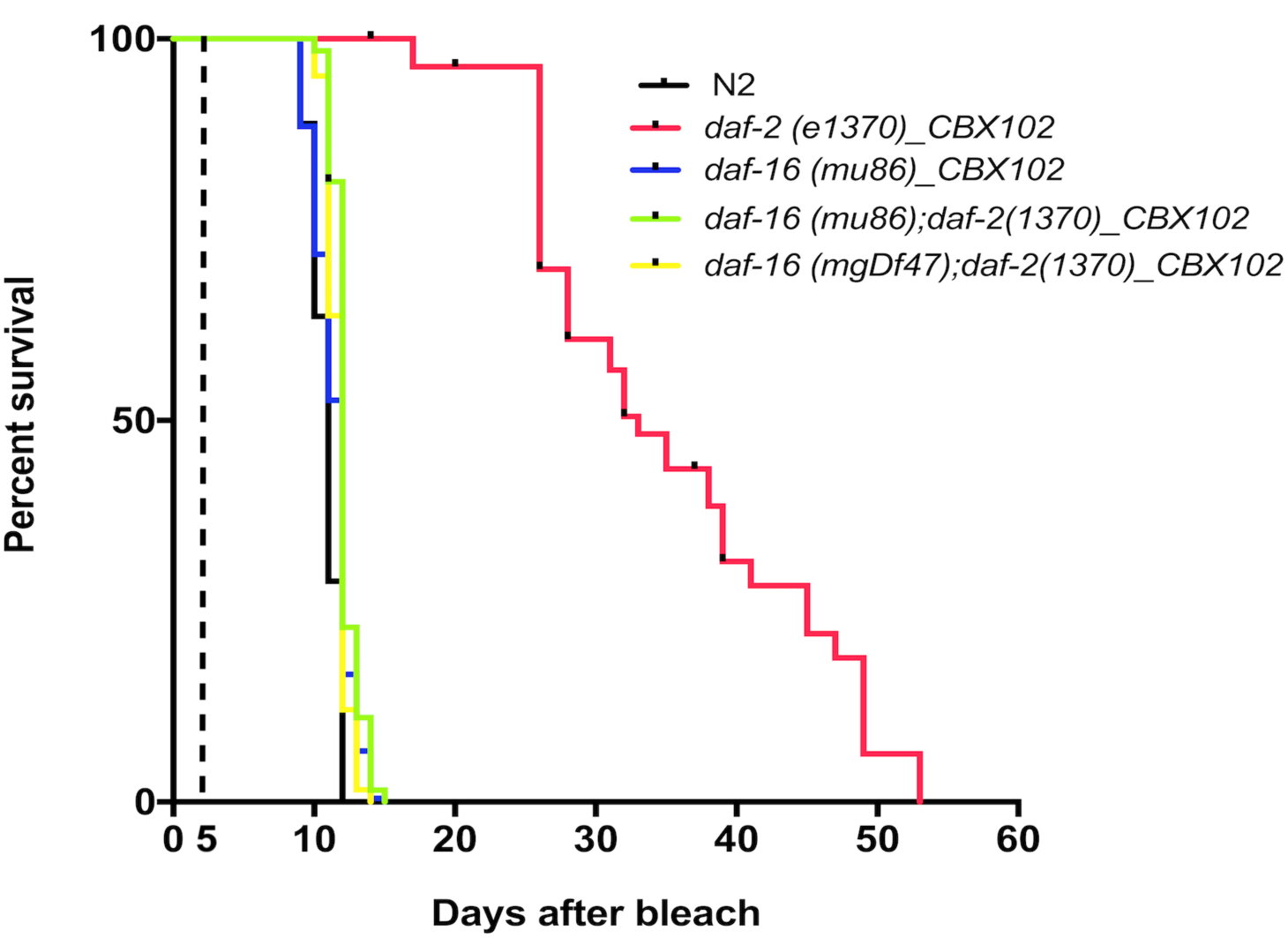
FOXO mediates the extension of *daf-2* lifespan on CBX102 under CGI. The *daf-2*-mediated lifespan extension on CBX102 was suppressed by *daf-16/FOXO*, using two mutants (*mu86* and *mgDf47*) of *daf-16*. We found that when compared to each other and to N2, both *daf-16, daf-2* double mutants as well as N2 had a lifespan with identical TD_50_ (12 days) on CBX102. This was also the lifespan TD_50_ of *daf-16(mu86)* alone (12 days). In contrast, lifespan of *daf-2* on CBX102 under CGI was significantly different (TD_50_=33, p<0.0001). For experiments involving the temperature sensitive *daf-2*, lifespan assays started at day 0 when animals were age-synchronized by bleach. Embryos were then left at 15^0^C on the appropriate bacterial diet till day 5. Day 5 marks the L4 to adult transition and time when plates were transferred to 25^0^C.

### *Daf-16* is required for the longevity and health of *daf-2* mutants under CGI

Life-span extension through the DAF-2 insulin-signalling pathway in *C. elegans* occurs by de-repression of the fork-head transcription factor DAF-16, which is normally under negative regulation by DAF-2. Therefore, strong loss-of-function alleles of *daf-16* such as *mgDf47* and *mu86* suppress the long-lived phenotype of *daf-2* under CGI with CBX102 (Fig 5). Moreover, *daf-16* exhibited a comparable degree of survival to CGI as N2 worms. Loss of DAF-16 suppressed the vigorous thrashing ability of *daf-2* making the double *daf-16; daf-2* statistically indistinguishable in its vigor compared to N2 (Fig S3). Therefore, the DAF-2/DAF-16 axis is important for maintaining longevity and health under CGI by a natural pathogen.

## DISCUSSION

Bacteria associated with the animal gut are important for gastrointestinal function (Fischbach 2018). Intestinal bacteria are involved in the synthesis and absorption of nutrients, protection of mucosal surfaces and the regulation of the immune function of the gut as well as influencing drug metabolism (Fischbach 2018). Quantitative and/or qualitative alterations of the intestinal microbiota underline many inflammatory diseases and chronic gastrointestinal infections (CGIs). In the short term, CGIs can lead to altered mucosal and immune function (Drossman *et al*, 2016). In the longer term, CGIs cause impaired epithelial barrier function (a major factor of reduced health span in old age) and changes in intestinal microbiota (dysbiosis) that can lead to constitutive inflammation in conditions like intestinal bowel disease and enterocolitis (Sperber and Dekel, 2010). We wanted to develop a simple model to test host longevity and health under CGI. *C. elegans* is such a model since microbiota=pathogen=food as the worm is a bacterial feeder and its laboratory culture is a mono-association.

Our work shows where longevity and immunity converge under CGI. Our data indicate that the insulin signalling pathway modulates both inherent longevity and pathogen resistance to affect overall survival across the life-course in a manner dependent on the pathogenicity of the bacteria on which *C. elegans* is feeding. The natural pathogen *M. nematophilum* strain CBX102 curtailed lifespan and health of N2 wild type worms but strain UV336 was statistically indistinguishable from *E. coli* OP50. Moreover, inactivating the insulin receptor via *daf-2* made worms live longer and be healthier and physiologically younger on CBX102. This correlated with reduced colonisation. In addition, UV336 extended *daf-2* lifespan even beyond what has been seen with *E. coli* OP50 acting as a probiotic when interacting with this host genetic background. More work is needed to identify the genetic differences between the two *M. nematophilum* strains and how lack of insulin host signalling modifies these bacterial strains and their properties.

The fact that inactivating the insulin pathway modifies a pathogen to become a commensal (in the case of CBX102) or a probiotic (in the case of UV336) may be evolutionarily conserved. Recent evidence in mice has shown that inducing insulin resistance through dietary iron drove conversion of a pathogen to a commensal. Specifically, insulin resistance converted the enteric pathogen *Citrobacter* to a commensal (Sanchez *et al*, 2018). There, reduced intestinal glucose absorbance was crucial for *Citrobacter* to be a commensal (Sanchez *et al*, 2018). More work is needed to determine if systemic glucose levels and/or intestinal glucose absorption play a role also in *C. elegans* and how this relates to the worm insulin pathway. However, reduced glucose levels increase lifespan (reviewed in Watts and Ristow, 2017). Reducing glycolysis has been shown to induce mitochondrial OXPHOS to generate a lifespan-extending reactive oxygen species (ROS) signal (Schulz *et al*, 2007) while increased levels have the opposite effect (Schulz *et al*, 2007; Zarse *et al*, 2012).

Taken together, our results and recent data from mice show that the consequences a microbe will cause to a host exist as a continuum. Thus, host genetics is important to determine where a microbe may lie in this continuum. The data show that the interaction between the worm and its bacterial food is a two-way interaction where host genes will play a role in shaping the long-term future of that interaction. In our system, the most prominent host proponent is the insulin-FOXO-dependent signalling pathway. *C. elegans* is an excellent model to design genetic screens and identify worm mutants that suppress the UV336-dependent extension of the *daf-2* longevity phenotype.

## MATERIAL AND METHODS

### *C. elegans* strains

All strains (supplementary table S1) were provided by the *Caenorhabditis* Genetic Center (CGC), University of Minnesota, and maintained at 20 °C, unless otherwise noted. The CGC is supported by the National Institutes of Health – Office of Research Infrastructure Programs (P40 OD010440).

### Bacteria growth conditions

*E. coli* OP50 or *M. nematophilum* (CBX102, UV336) cultures were grown in LB at 37 °C. Bacterial lawns were prepared by spreading 100 μl of an overnight culture on a 6 cm diameter NGM plate. Plates were incubated overnight at room temperature.

### Immunity and longevity Assays

CBX102 assays were performed at 25 °C, unless otherwise noted, as previously described (Gravato-Nobre *et al*. 2016, Plos Pathogens). To test/validate immunity or longevity phenotypes of *daf-2* (e1370), worms were raised on CBX102 or OP50 to the L4 stage at the permissive temperature (15 °C), and shifted to the restrictive temperature of 25 °C.Worms were age-synchronized by bleaching and embryos were incubated at 25 °C on NGM agar plates with lawns of *E. coli* OP50 or *M. nematophilum* CBX102. The embryonic stage (day of bleach) was designated as Day 0. A total of 125 worms were used per lifespan assay. On day 2, 25 animals were transferred to each NGM plate. Animals were scored daily and transferred to fresh lawns every other day. Death was defined when an animal no longer responded to touch. Worms that died of bagging or crawled off the plates were censored from the analysis. For each mutant population and bacterial lawn, the time required for 50% of the animal to die (TD50) was compared to that of the control populations using a *t* test. A *p-*value*<* 0.05 was considered significantly different from the control.

### SYTO 13 staining

Overnight bacterial cultures were concentrated 10x by spinning them at 2500 rpm, and their pellet suspended in 1 ml of TBS containing 3 μl of SYTO 13. Bacterial colonization was determined by exposing the animals to SYTO13-labelled CBX102 or OP50. To allow for their complete post-embryonic development, animals were left on CBX102 lawns, at 15 °C until most mutant animals reached L4, after swhich they were shifted to 25 °C for another day. On day 7, one-day-old adult worms were exposed to SYTO 13-labelled CBX102. Worms were visualized after 20 hours of feeding on SYTO 13-labeled CBX102. Live worms were mounted on a glass slide in 25 μM tetramisole on a 2% agarose pad and examined using a Leica SP5 confocal microscope.

### Thrashing Assays

One-day old adults were placed in a drop of M9 and allowed to recover for 40 s (to avoid behaviour associated with stress), after which animal were video recorded for 30s. The number of body bend per second (BBPS) was determined by importing captured video images to ImageJ and by using wrMTrck plugin developed by Jesper S, Pederson. (http://www.phage.dk/plugins/wrmtrck.html). More than 20 animals were used in each treatment. Thrashing experiments were done in triplicates. All statistical analysis data performed using GraphPad Prism software.

## ACKNOWLEDGMENTS

This work was funded by the EP Abraham Cephalosporin Trust grant no. CF 319 (to PL).

## FIGURE LEGENDS

**FIGURE S1.**
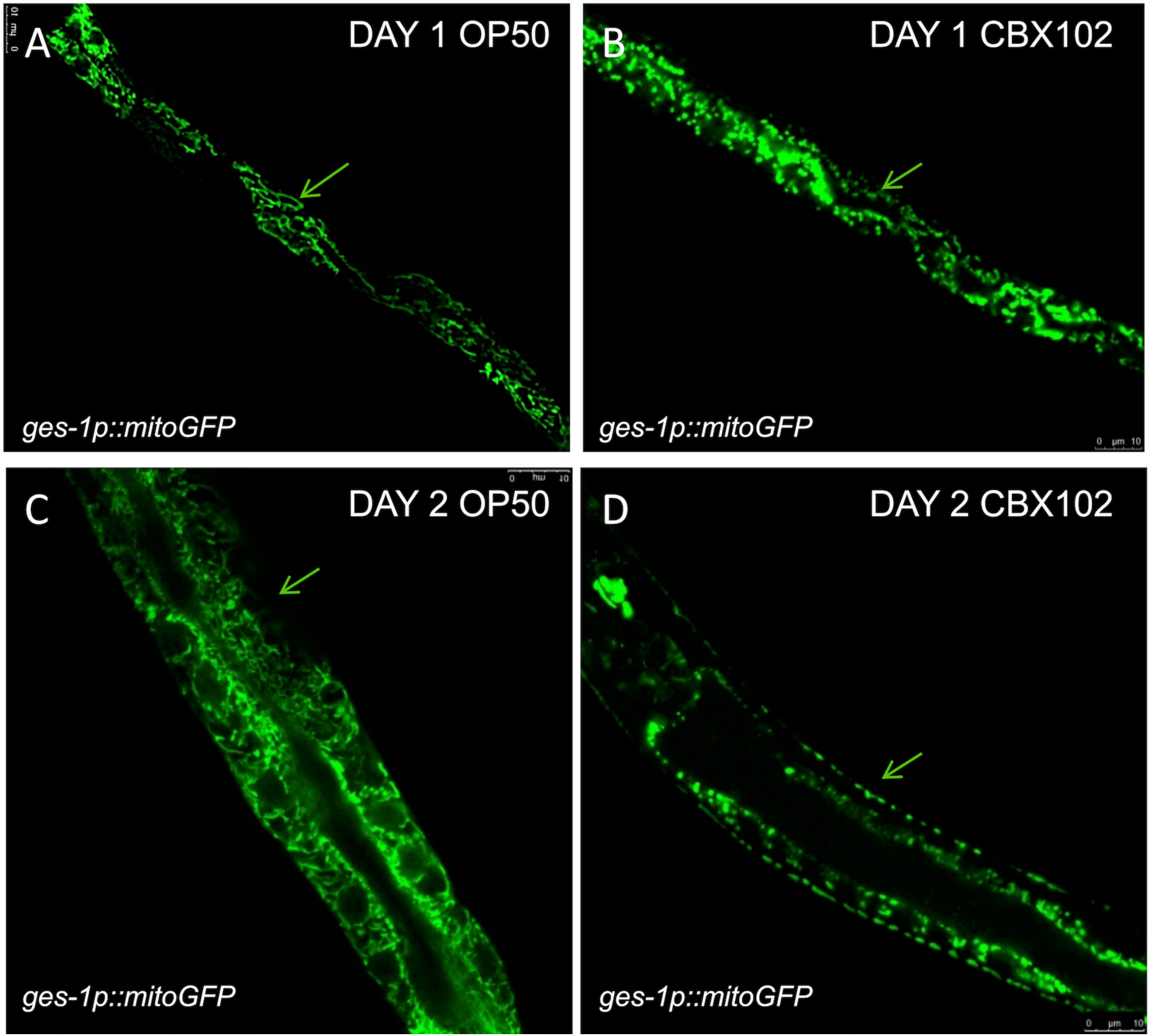
*M. nematophilum* CBX102 accelerates ageing. Animals expressing the mitochondria marker *mito-GFP* in the intestine **(A), (C)** in OP50 showing normal tubular mitochondria while age-matched **(B), (D)** CBX102-grown L2 animals show fragmented mitochondria with irregular shape.

**FIGURE S2.**
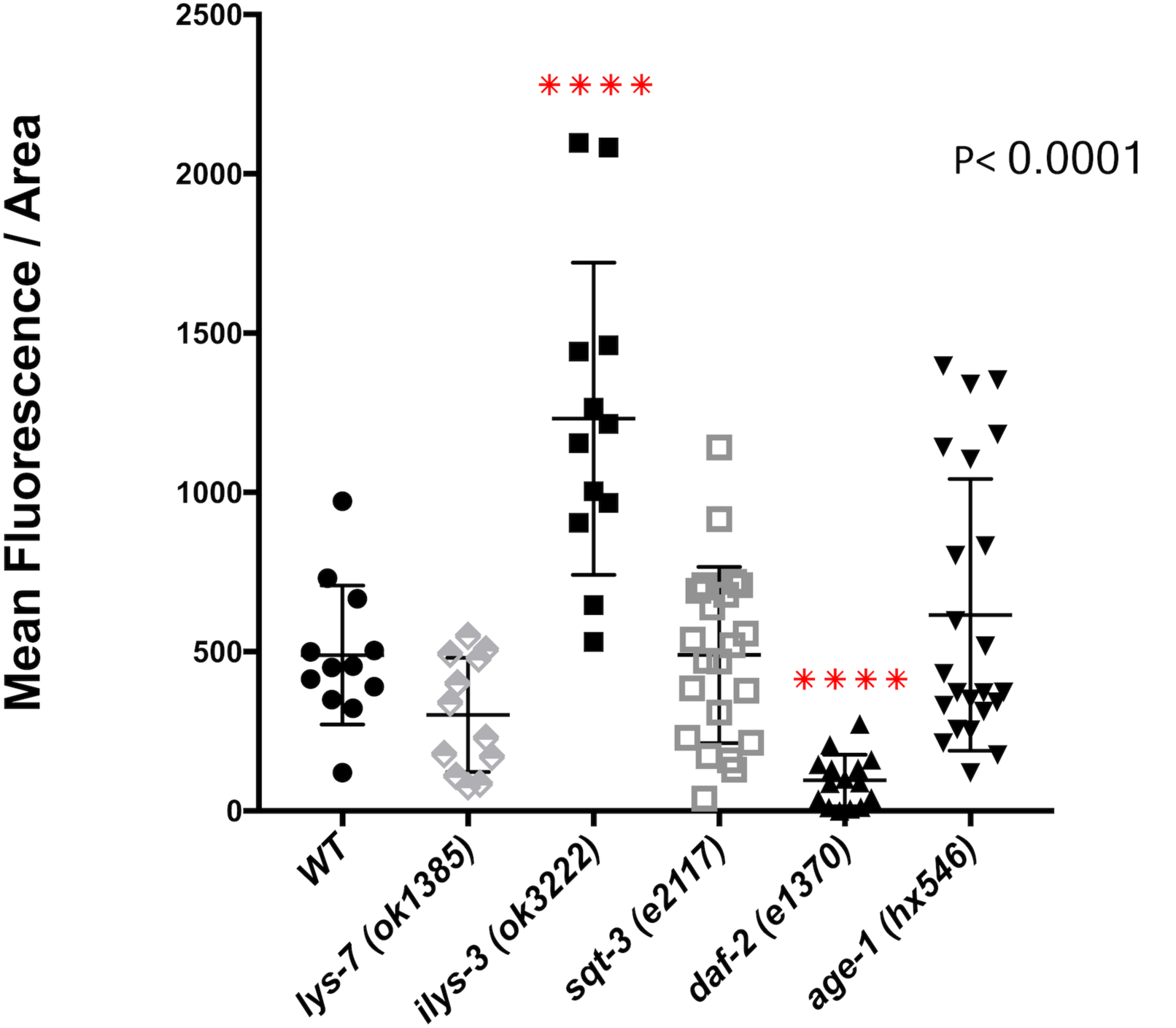
Bacterial colonisation of CBX102 in a *C. elegans* mutants. Each dot represents a 1-day old animal with SYTO13 fluorescence counted. *M. nematophilum* strain CBX102 displayed less colonisation in *daf-2* compared to N2 (designated as wild-type of WT). In contrast, mutants lacking the antimicrobial *ilys-3* gene, displayed significantly increased colonisation. Dunnett’s-multiple comparisons one-way ANOVA test was performed. \*\*\*\**P*<0.0001; except comparisons with *ilys-3* and *daf-2*, all other comparisons were not significant.

**FIGURE S3.**
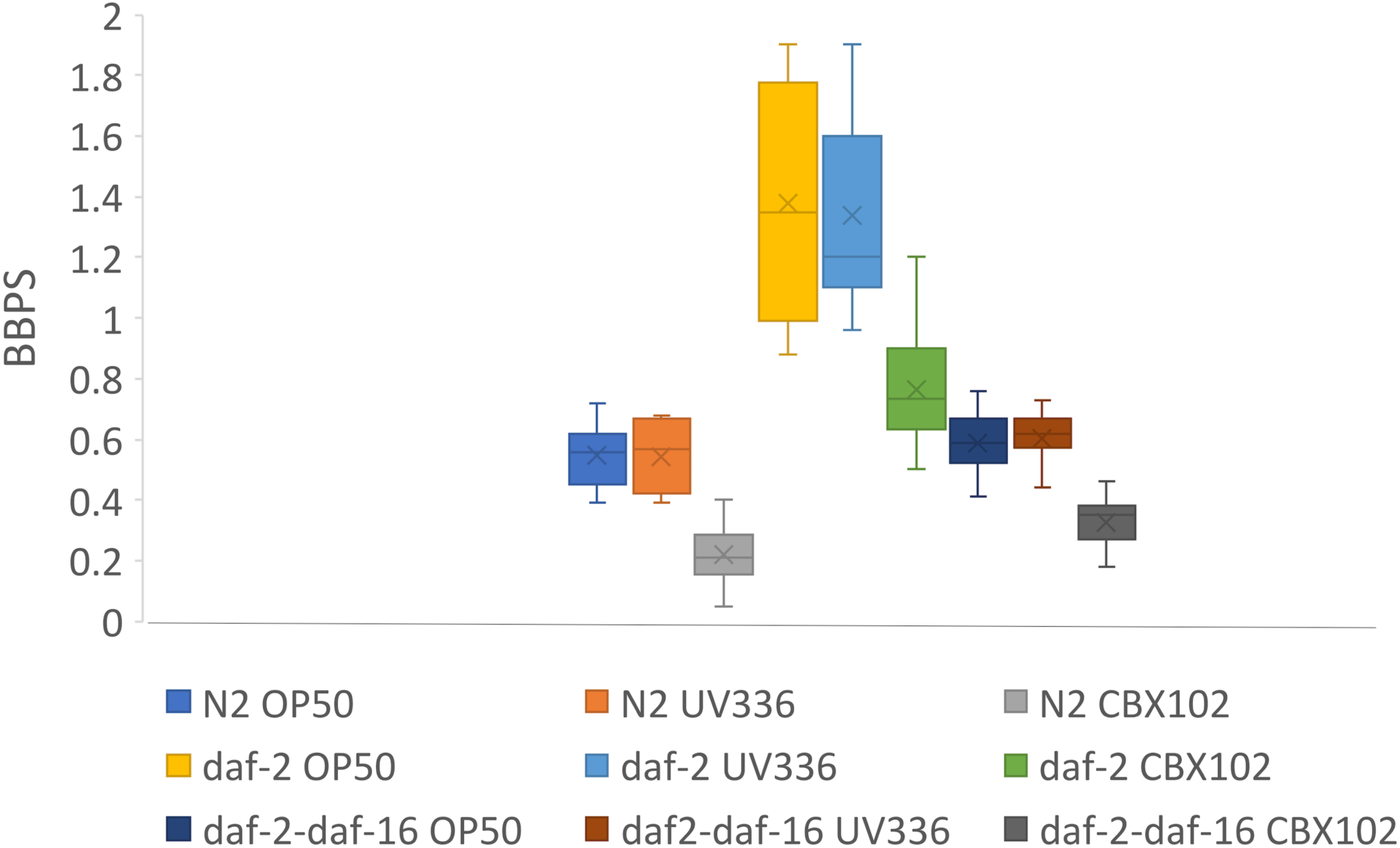
Health of animals with different microbiota. Box plot represents body bends per second (BBPS) counted per animal for each strain. Each box represents a group of 1-day old animals (n=25). Dunnett’s-multiple comparisons one-way ANOVA test was performed showing that CBX102 was always significantly lower across the same host genotypes (p<0.0001) while *daf-2* was significantly higher than N2 across bacterial strains (p<0.0001). Comparison between *daf2* and *daf-16, daf-2* showed significant difference (p<0.0001) across the different bacteria while N2 and *daf-16, daf-2* were statistically indistinguishable (p>0.1).

**Table S1.**
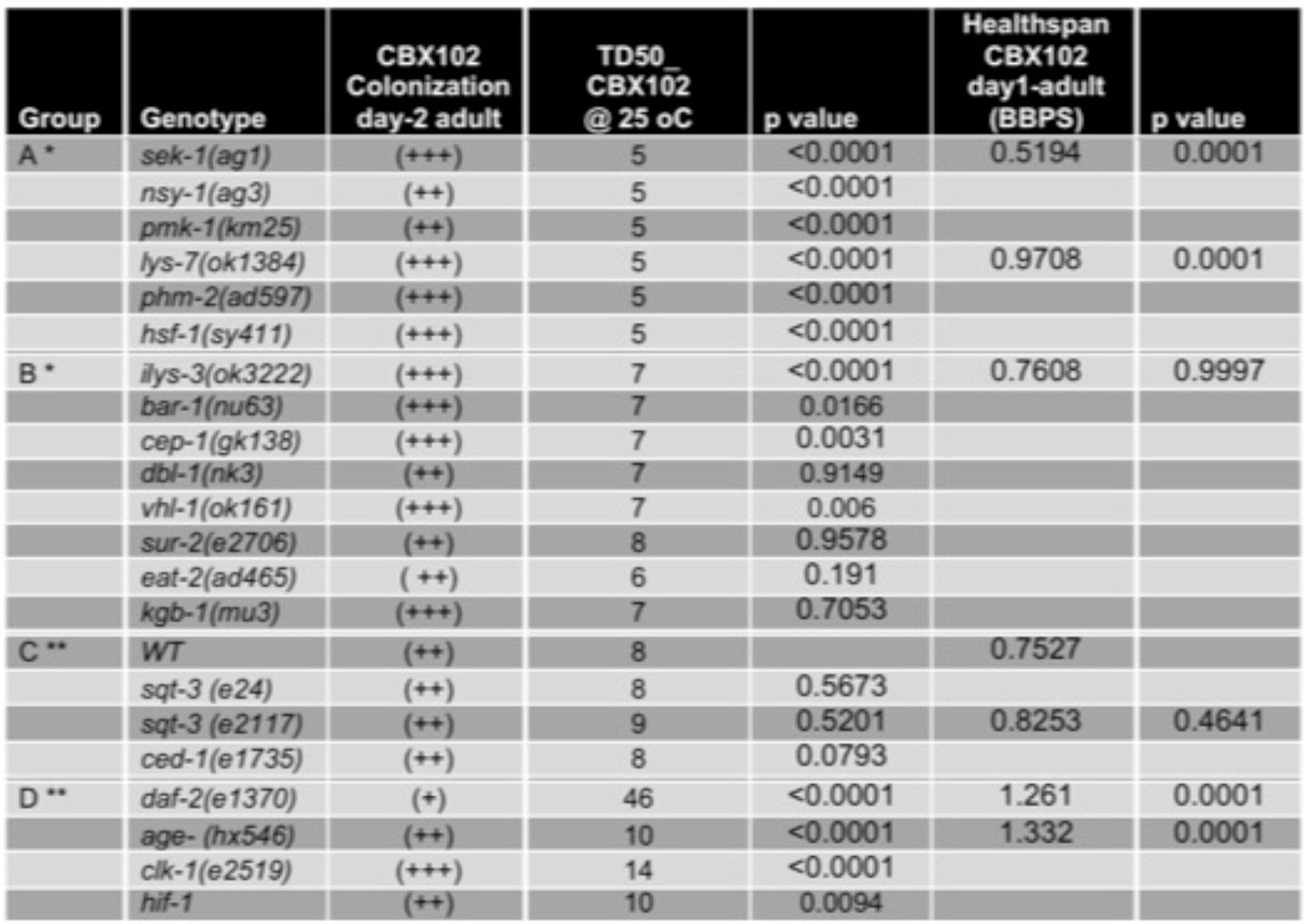
Statistics for Lifespan and Health span assays and mutants tested. For group categories see Fig. 2. WT is wild type (strain N2). Measurements: *relative to *ilys-3* and **relative to WT (N2). *C. elegans* mutants without a numerical value in the health span column were not moving at all and therefore we were unable to film their vigour.

